# The properties of 5-methyltetrahydrofolate dehydrogenase (MthfD) and its role in the tetrahydrofolate (THF)-dependent dicamba demethylation system in *Rhizorhabdus dicambivorans* Ndbn-20

**DOI:** 10.1101/537639

**Authors:** Shigang Yao, Le Chen, Zhou Yang, Li Yao, Jianchun Zhu, Jiguo Qiu, Guoxiang Wang, Jian He

**Author notes:** Address correspondence to Jian He.

## Abstract

The herbicide dicamba is initially degrade via demethylation in *Rhizorhabdus dicambivorans* Ndbn-20. A gene cluster scaffold 66 containing a THF-dependent dicamba methyltransferase *dmt* and three THF metabolic-related genes, namely, *mthfD*, *dhc* and *purU*, is responsible for dicamba demethylation in this strain. However, the characteristics and functions of MthfD, Dhc and PurU have not been elucidated. In this study, MthfD was synthesized in *Escherichia coli* BL21(DE3) and purified as a His_6_-tagged protein. Purified MthfD was found to be a monomer, and exhibited 5-CH_3_-THF dehydrogenase activity *in vitro*. The *K*_cat_ and *K*_m_ for 5-CH_3_-THF were 0.23 s^−1^ and 16.48 μM, respectively. However, 5,10-CH_2_-THF reductase activity was not detected for MthfD yet. Gene disruption results showed that *mthfD* is essential for dicamba degradation, whereas *dhc* is dispensable. Our studies revealed that MthfD physiologically is a 5-CH_3_-THF dehydrogenase that catalyzes the irreversible dehydrogenation of 5-CH_3_-THF to 5,10-CH_2_-THF in the THF regeneration pathway during dicamba demethylation in *R. dicambivorans* Ndbn-20.

## IMPORTANCE

The THF-dependent demethylase system is an important demethylation mechanism that is responsible for demethylation of many natural and synthetic methyl-substituted compounds. However, the mechanism underlying THF regeneration and the characteristics and function of enzymes involved have not been elucidated. *mthfD* encodes a protein that shares the highest sequence identity with MetF, which physiologically acts as a 5,10-CH_2_-THF reductase in organisms. This study investigated the characteristics, catalytic activities and function of MthfD. The results revealed that MthfD differs from previously reported MetF in phylogenesis, characteristics and function. MthfD physiologically acts as a 5-CH_3_-THF dehydrogenase but not 5,10-CH_2_-THF reductase, and it is essential for dicamba catabolism in *R. dicambivorans* Ndbn-20. This study provides new insights into the mechanism of THF-dependent methyltransferase system.

## INTRODUCTION

Methyl-substituted compounds such as caffeine, chloromethane, sesamin, vanillate and syringate, are widely present in nature (1–6). And a variety of synthetic methyl-substituted aromatics are used in industrial and agricultural production, e.g., dicamba, linuron, chlortoluron, alachlor and isoproturon, which are important herbicides ((7–10). Demethylation is usually the initial and critical step in the microbial degradation of methyl-substituted compounds. To date, three types of demethylases from aerobic bacteria have been reported: Rieske non-heme iron oxygenase (RHO) (11, 12), cytochrome P450 (13, 14) and tetrahydrofolate (THF)-dependent methyltransferase (6, 15–21). Both RHO and P450 are multi-component monooxygenase systems with NAD(P)H as cofactor, these demethylases add a molecule of oxygen to the methyl group, resulting in removal of the methyl group from the substrate to form formaldehyde. THF-dependent methyltransferase transfers the methyl from the substrates to cofactor THF producing 5-methyltetrahydrofolate (5-CH_3_-THF), thus its process and mechanism are completely different from RHO- and P450-mediated demethylation. To date, five THF-dependent methyltransferases have been identified: the vanillate methyltransferase LigM and syringate methyltransferase DesA from *Sphingomonas paucimobilis* SYK-6 (6, 17), the chloromethane methyltransferase CmuAB from *Methylobacterium chloromethanicum* CM4 (18, 20, 21) and the dicamba methyltransferases Dmt and Dmt50 from *Rhizorhabdus dicambivorans* Ndbn-20. (16, 19).

Regeneration of THF from 5-CH_3_-THF is a vital step in the THF-dependent demethylation system. In *S. paucimobilis* SYK-6, two THF metabolism-related genes, namely, *metF* [putative 5,10-methylenetetrahydrofolate (5,10-CH_2_-THF) reductase gene] and *ligH* (putative formyl-THF deformylase gene, also named *purU*) are located downstream of *ligM* (17). In *R. dicambivorans* Ndbn-20, two dicamba methyltransferase gene clusters, namely, scaffold 66 and scaffold 50, were identified, and each contains three THF metabolism-related genes, namely, *metF*, *dhc* (putative 5,10-CH_2_-THF dehydrogenase/5,10-methenyl-THF cyclohydrolase gene) and *purU* (16, 19, Fig. 1A). And in *M. chloromethanicum* CM4, the chloromethane methyltransferase gene cluster also contains three THF metabolism-related genes, *metF, fold* (*dhc*) and *purU* (20). Based on the functional prediction of these genes, a possible pathway for THF regeneration from 5-CH_3_-THF was proposed: 5-methyl-THF dehydrogenates to 5,10-CH_2_-THF under the catalysis of MetF, 5,10-CH_2_-THF further dehydrogenates to 5,10-methenyltetrahydrofolate (5,10-CH=THF) and then is cyclohydrolased to 10-CHO-THF by bifunctional enzyme Dhc (FolD), and finally, 10-CHO-THF is deformed to THF by PurU (LigH) (6, 19, 21, Fig. 1C). However, this THF cycling pathway has not been validated by enzymatic studies, and none of the three THF metabolism-related enzymes involved has been purified, functionally identified or characterized.

**FIG 1.**
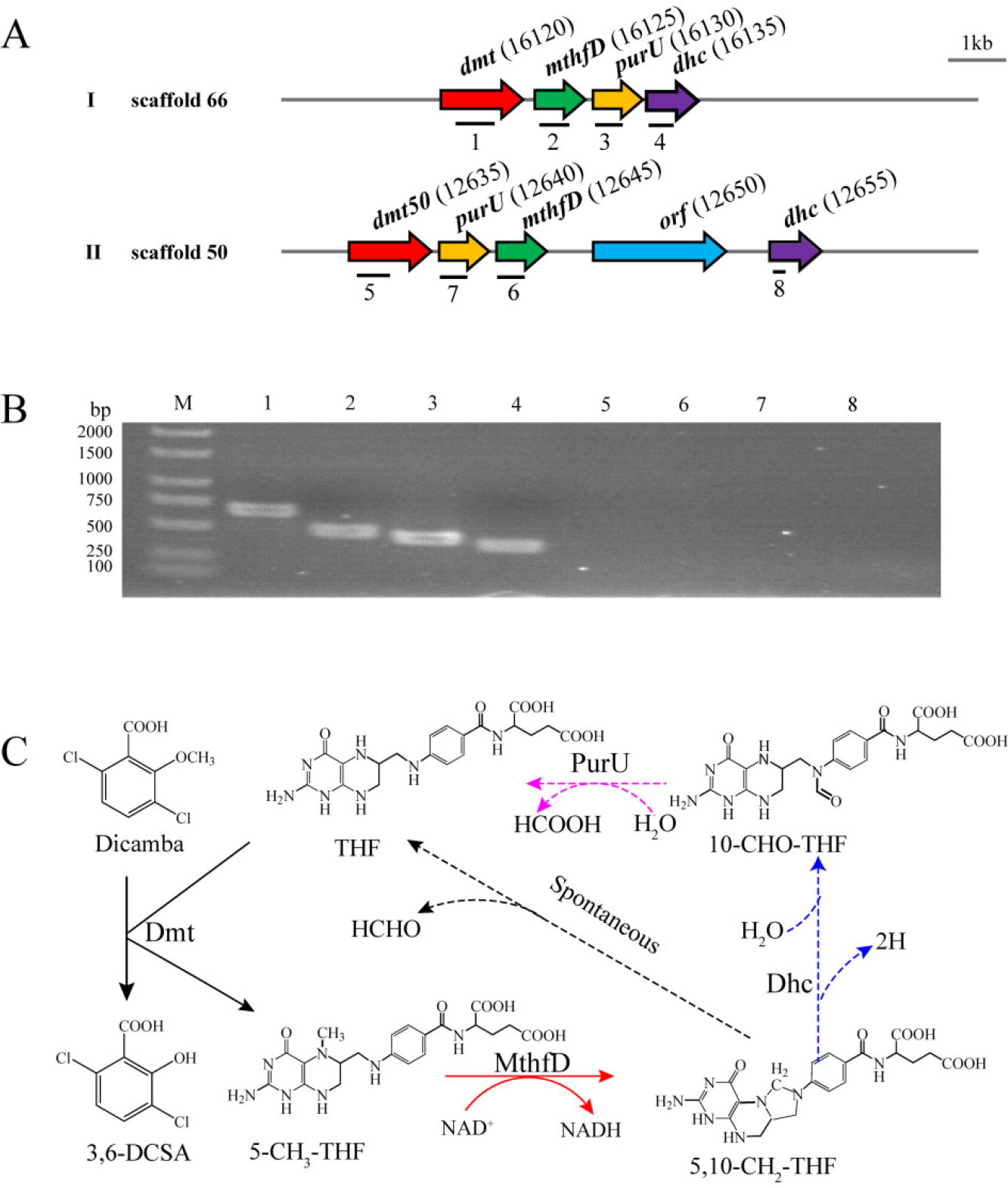
(A) Organization of the THF-dependent dicamba methyltransferase gene clusters scaffold 66 and scaffold 50 in the genome of *R. dicambivorans* Ndbn-20. *dmt and dmt50*, dicamba methyltransferase gene; *mthfD*, 5-CH_3_-THF dehydrogenase gene; *dhc*, 5,10-CH_2_-THF dehydrogenase/5,10-methenyl-THF cyclohydrolase gene; *purU*, formyl-THF deformylase gene. Number in parentheses indicates the locus tag for each gene in the genome. Arrow indicates the size and transcriptional direction of each gene. Lines below the gene cluster show the locations and sizes of the PCR fragments in panel B. (B) Transcription analysis of *dmt*, *dmt50*, *mthfD*, *purU* and *dhc* by RT-PCR. (C) Proposed regeneration pathway of THF from 5-CH_3_-THF in *R. dicambivorans* Ndbn-20. The reactions catalyzed by Dhc and PurU have not been confirmed by enzymatic studies.

MetF (also referred to as MTHFR) is the rate-limiting enzyme in C1 metabolism in organisms (22–24). MetF physiologically catalyzes the reduction of 5,10-CH_2_-THF to 5-CH_3_-THF (5,10-CH_2_-THF reductase activity) using flavinadenine dinucleotide (FAD) as a prosthetic group (25). MetF also catalyzes the reverse reaction (5-CH_3_-THF dehydrogenase activity) *in vitro* (26). To date, many studies have reported the function and characteristics of MetF from *E. coli*, *Moorella thermoacetica*, yeast, *Arabidopsis thaliana* and horse. Eukaryotic MetF is a homodimer composed of 70-77 kDa subunits, each of which has a N-terminal catalytic domain and a C-terminal regulatory domain (27). In mammals and yeasts, MetF uses NADPH as electron donor, and the reduction reaction catalyzed by these MetF is physiologically irreversible and is inhibited by S-adenosylmethionine (AdoMet) (27). The plant MetF uses NADH as electron donor and does not contain a regulatory domain, and the reaction catalyzed by plant MetF is physiologically reversible (28). The prokaryotic MetF subunit is approximately 30 kDa in size; the MetF from *E. coli* is a tetramer (26), while the MetF from *M. thermoacetica* is a hexamer (29). Similar to plant MetF, prokaryotic MetF also uses NADH as electron donor and has no regulatory domain (26).

In this study, the *metF*-like ORF in the dicamba methyltransferase gene cluster scaffold 66 from *R. dicambivorans* Ndbn-20 was renamed as *mthfD* (encoding 5-<u>m</u>ethyl<u>t</u>etra<u>h</u>ydro<u>f</u>olate <u>d</u>ehydrogenase); *mthfD* was overexpressed in *E. coli* BL21(DE3), and the His_6_-tagged MthfD was purified using Co-affinity chromatography. The characteristics and physiological function of this enzyme in the THF regeneration during dicamba demethylation in *R. dicambivorans* Ndbn-20 were investigated.

## RESULTS

### Expression of *mthfD* and purification of the recombinant MthfD

To investigate the catalytic activity and characteristics of MthfD, *mthfD* and *metF* of *E. coli* were expressed in *E. coli* BL21(DE3) using the pET29a(+) expression system, and the C-terminal His_6_-tagged proteins were purified to apparent homogeneity using Co^2+^-chelate affinity chromatography. Sodium dodecyl sulfate (SDS)-polyacrylamide gel electrophoresis (PAGE) analysis showed that the molecular weight of both MthfD and *E. coli* MetF were approximately 32 kDa (Fig. S1), which was consistent with the deduced molecular weights (31.8 kDa and 33.1 kDa, respectively). Gel filtration showed that the molecular mass of the native MthfD and *E. coli* MetF were 34.2 kDa and 110.2 kDa, respectively (Fig. 2), indicating that the native MthfD is a monomer, and the *E. coli* MetF is a tetramer. In previous study, the *E. coli* MetF was also identified as a homotetramer (26). The purified MthfD and *E. coli* MetF appeared to be yellow because the two proteins contained flavin adenine dinucleotide (FAD) as a prosthetic group (30). MthfD remained yellow after long-term dialysis against Tris-HCl buffer or heat treatment in a boiling water bath, indicating that FAD tightly binds to MthfD; whereas *E. coli* MetF was found to easily lose FAD upon dilution. When stored at − 80°C for two months in 20 mM Tris-HCl buffer with 10% glycerol and 0.3 mM EDTA, 90% activity of MthfD was retained, indicating that MthfD was stable. In contrast, the *E. coli* MetF was prone to lose enzymatic activity, only 30% of its activity was retained when stored at −80°C for two months.

**FIG 2.**
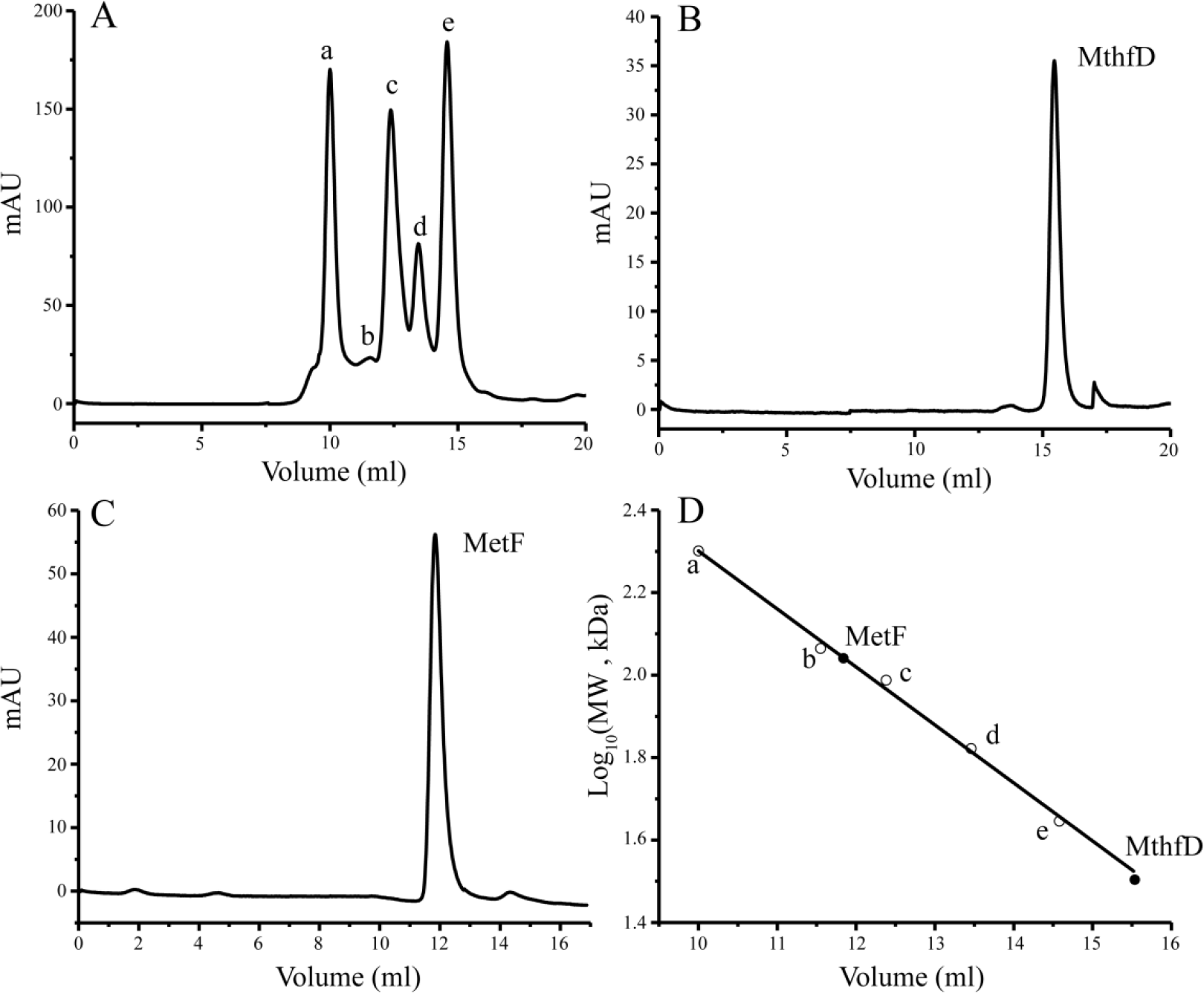
Gel filtration of the purified MthfD and *E. coli* MetF. (A) The peak volumes of standard proteins: a, myosin (200.0 kDa,10.00 mL); b, β-galactosidase (116.0 kDa, 11.55 mL); c, phosphorylase b (97.2 kDa, 12.38 mL); d, bovine serum albumin (66.4 kDa, 13.46 mL); e, ovalbumin (44.3 kDa, 14.58 mL); (B) The peak volume of native MthfD (15.52 mL); (C) The peak volume of native *E. coli* MetF (11.84 mL); (D) The calibration line of standard proteins and the molecular mass determination of MthfD (33.4 kDa) and *E. coli* MetF (110.2 kDa).

### 5-CH_3_-THF dehydrogenase activity

The physiological function of MthfD was predicted to catalyze the dehydrogenation of 5-CH_3_-THF, generated during dicamba demethylation, to 5,10-CH_2_-THF in *R. dicambivorans* Ndbn-20 (Fig. 1C). According to the enzymatic assay, both MthfD and *E. coli* MetF exhibited 5-CH_3_-THF dehydrogenase activities in the presence of menadione. The product 5,10-CH_2_-THF was unstable and spontaneously decomposes to THF and formaldehyde, as identified by high-performance liquid chromatography (HPLC) (Fig. 3). The spontaneous decomposition of 5,10-CH_2_-THF has also been observed in previous studies (31). The kinetic parameters of MthfD for 5-CH_3_-THF were determined as follows: the *K*_cat_ was 0.23 ± 0.01 s^−1^, the *K*_m_ was 16.48 ± 1.81 μM (Fig. S2A), and the *K*_cat_/*K*_m_ was 14.17 s^−1^ mM^−1^. In contrast, for *E. coli* MetF, the *K*_cat_ was 0.16 ± 0.01 s^−1^, the *K*_m_ was 30.95 ± 3.77 μM (Fig. S2B), and the *K*_cat_/*K*_m_ was 5.17 s^−1^ mM^−1^. Obviously, MthfD showed higher affinity and catalytic efficiency toward 5-CH_3_-THF than that of *E. coli* MetF.

**FIG 3.**
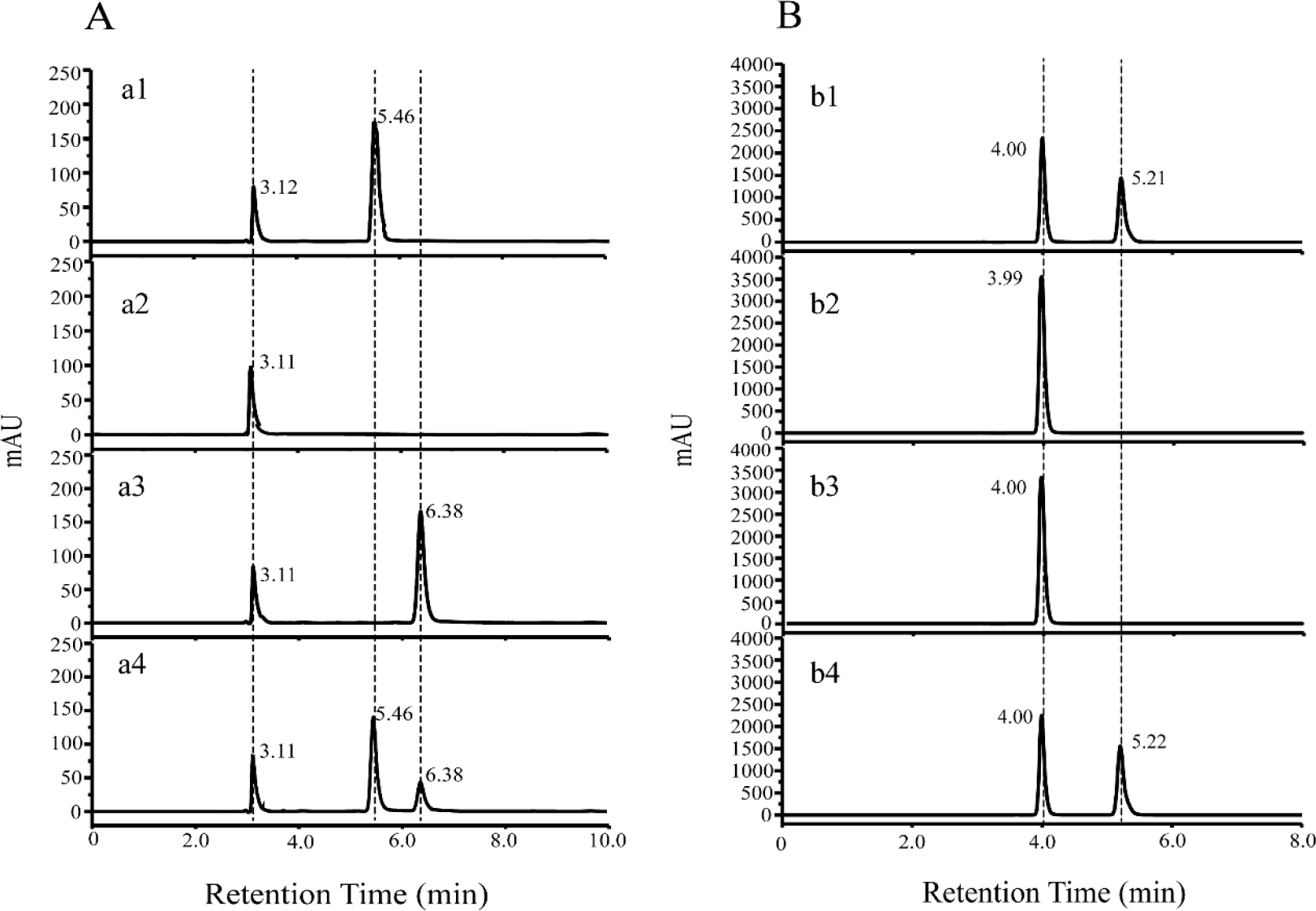
HPLC analysis of the product generated during 5-CH_3_-THF dehydrogenation by MthfD. (A) HPLC analysis of the product THF. a1, THF standard; a2, ascorbic acid standard; a3, 5-CH_3_-THF control without addition of enzyme; a4, product generated during 5-CH_3_-THF dehydrogenation by MthfD. 5.46 min peak represents THF, 3.11 min peak represents ascorbic acid added in the solvent or enzymatic mixture to maintain a reducing environment, 6.38 min peak represents 5-CH_3_-THF. (B), HPLC analysis of the product formaldehyde. b1, formaldehyde standard; b2, 2,4-dinitrophenylhydrazine standard; b3, Control added with 5-CH_3_-THF but no enzyme; b4, product generated during 5-CH_3_-THF dehydrogenation by MthfD. 5.21 min peak represents 2,4-dinitrobenzenehydrazone, 4.00 min peak represents 2,4-dinitrophenylhydrazine.

### 5,10-CH_2_-THF reductase activity

In previous reports, the physiological reaction catalyzed by MetF is the reduction of 5,10-CH_2_-THF to 5-CH_3_-THF (24–28). To determine whether MthfD possesses 5,10-CH_2_-THF reductase activity, 5,10-CH_2_-THF was prepared by spontaneous reaction of THF and excess formaldehyde, as 5,10-CH_2_-THF is unstable. The 5,10-CH_2_-THF reductase activity was determined by monitoring the disappearance of NADH, which has characteristic absorption peak at 340 nm. The result in Fig. 4A shows a rapid decrease in the absorption peak at 340 nm in the enzymatic mixture added with *E. coli* MetF, indicating that *E. coli* MetF exhibited the 5,10-CH_2_-THF reductase activity. However, the absorption peak at 340 nm in the enzymatic mixture added with MthfD remained unchanged (Fig. 4B), indicating that MthfD could not reduce 5,10-CH_2_-THF to 5-CH_3_-THF or the 5,10-CH_2_-THF reductase activity was very low.

**FIG 4.**
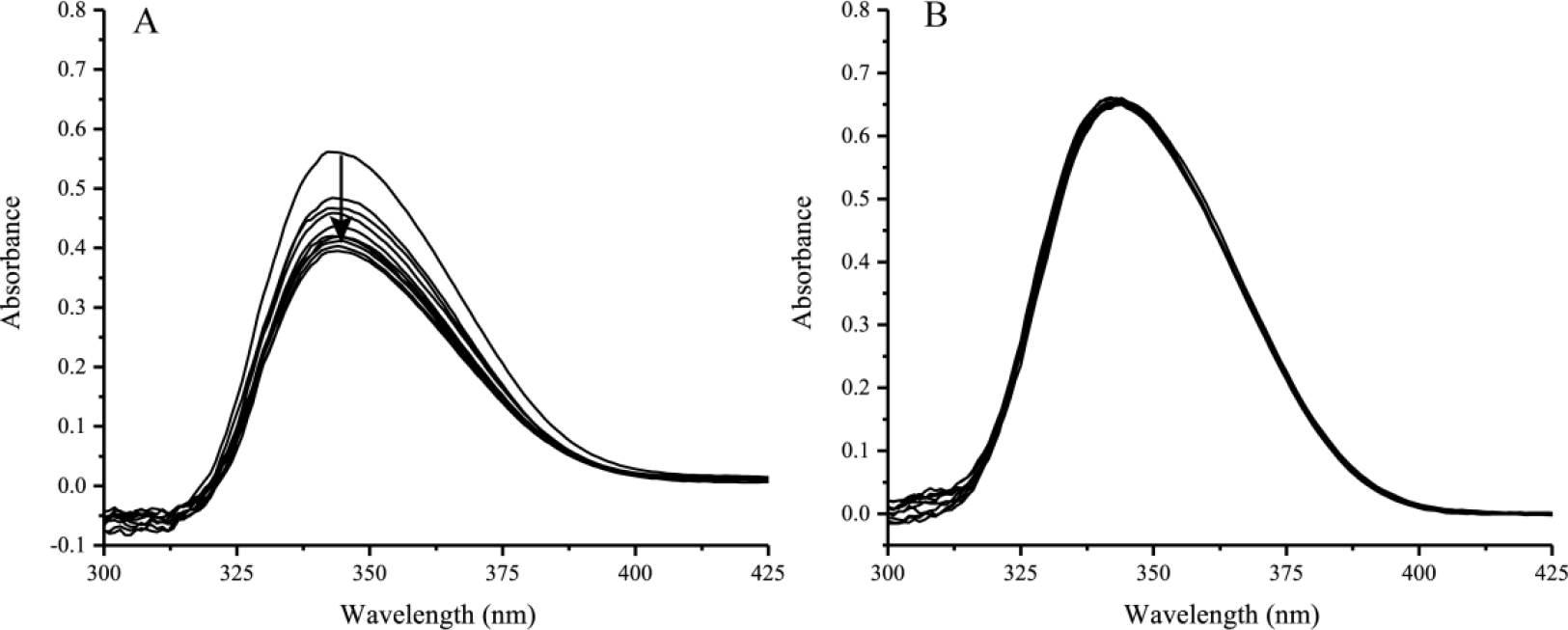
Spectrophotometric changes during the determination of 5,10-CH_2_-THF reductase activities of *E. coli* MetF (A) and MthfD (B). The spectra were recorded every 30 s. The arrow indicates the direction of spectral changes.

### Transcriptional analysis of *mthfD* in *R. dicambivorans* Ndbn-20

In our previous study, two THF-dependent dicamba methyltransferase gene clusters (scaffold 66 and scaffold 50) was identified in *R. dicambivorans* Ndbn-20. Each cluster included a dicamba methyltransferase gene (*dmt* or *dmt50*) and three THF metabolism-related genes, namely, *mthfD*, *purU* and *dhc* (16, 19). To investigate whether the genes in the two clusters were transcribed in dicamba-cultured cells of *R. dicambivorans* Ndbn-20, reverse transcription-PCR (RT-PCR) was performed using RNA extracted from cells grown on dicamba. The results showed that *dmt*, *mthfD*, *purU* and *dhc* in scaffold 66 were transcribed, whereas the genes in scaffold 50 were not transcribed, indicating that genes in cluster scaffold 66 was responsible for dicamba demethylation in *R. dicambivorans* Ndbn-20 (Fig. 1B).

### Disruption of *mthfD* and *dhc* from *R. dicambivorans* Ndbn-20

To investigate the physiological role of MthfD in dicamba demethylation in *R. dicambivorans* Ndbn-20, the genes *mthfD* and *dhc* were individually disrupted by homologous recombination. Two mutants, namely, Ndbn-20∆*mthfD* and Ndbn-20∆*dhc*, were generated. There was no significant difference in growth rate and morphology between the two mutants and wild-type when the cells were cultured in 1/5 Luria-Bertani (LB) broth or 1/5 LB agar (data not shown). The results of the whole-cell transformation experiments showed that mutant Ndbn-20∆*mthfD* entirely lost the ability to degrade dicamba and utilize dicamba (Fig. 5A). It is interesting that like the wild-type (Fig. 5B), mutant Ndbn-20∆*dhc* degraded 98.9% of the added 1.0 mM dicamba within 36 h of incubation, and the OD_600 nm_ of the cultures increased from 0.20 to 0.39 (Fig. 5C). The above results indicated that *mthfD* is essential for the degradation and utilization of dicamba by strain Ndbn-20, whereas *dhc* is dispensable for dicamba metabolism.

**FIG 5.**
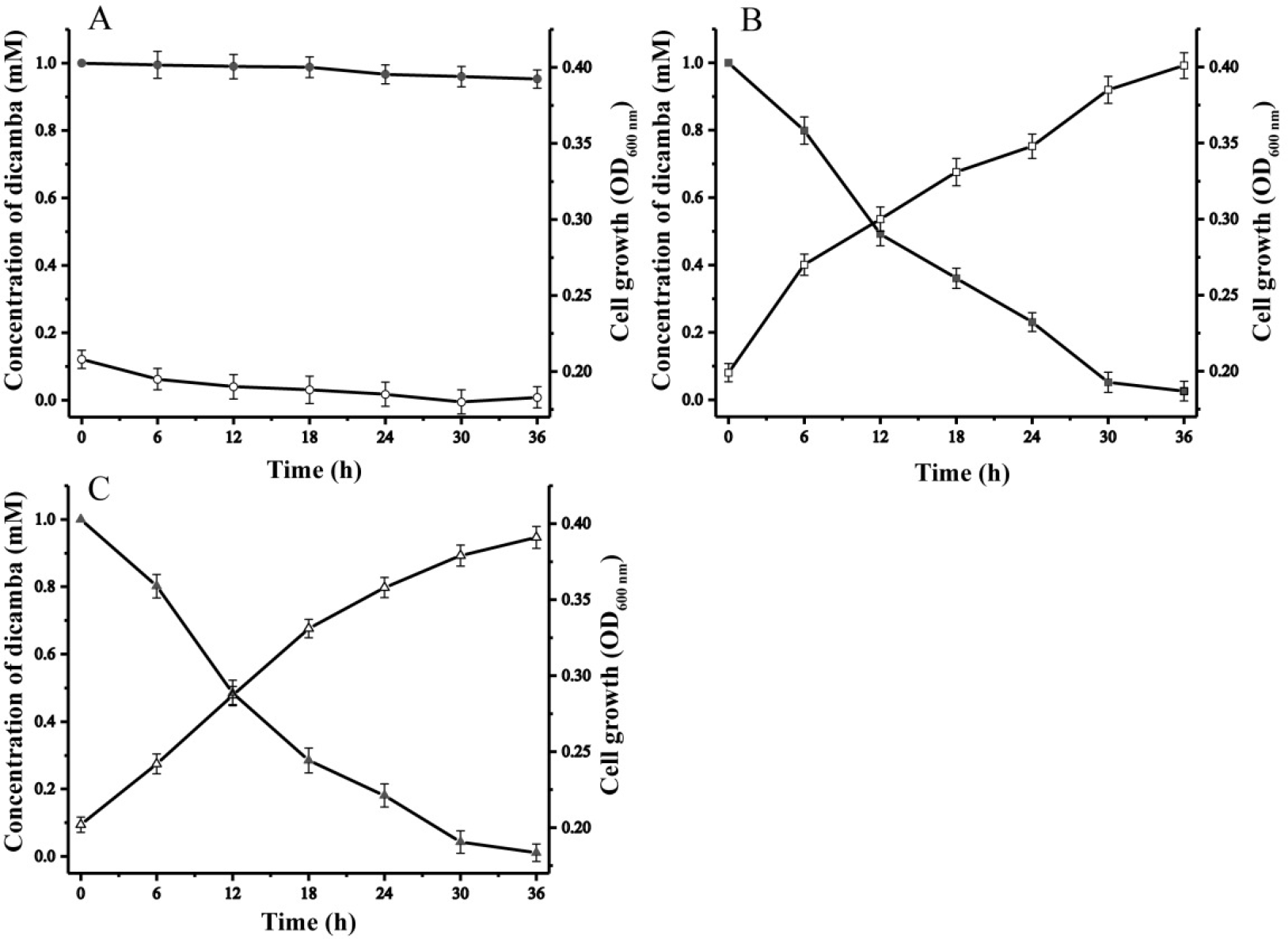
Time course of dicamba degradation by mutant Ndbn-20∆*mthfD* (A), wild-type (B) and mutant Ndbn-20∆*dhc* (C). The data were derived from three independent measurements, and the error bars indicate standard deviations.

### 5-CH_3_-THF dehydrogenation by the cell lysates of *R. dicambivorans* Ndbn-20 and the *mthfD*- or *dhc*-disrupted mutants

To further study the physiological function of MthfD in THF regeneration in *R. dicambivorans* Ndbn-20, the cell lysates of the mutants Ndbn-20∆*mthfD* and Ndbn- 20∆*dhc* and the wild-type were prepared from cells grown on dicamba and harvested in the logarithmic phase. The enzymatic study showed that the cell lysate of mutant Ndbn-20∆*mthfD* completely lost the 5-CH_3_-THF dehydrogenase activity (Fig. S3A). The results further confirmed that MthfD was the only active 5-CH_3_-THF dehydrogenase in cells grown on dicamba. Furthermore, although the *dhc* was disrupted in mutant Ndbn-20∆*dhc*, its cell lysate could still transform 5-CH_3_-THF to THF at a rate similar to that of the wild-type. (Fig. S3B).

### Regeneration of THF by purified MthfD from 5-CH_3_-THF produced during dicamba demethylation

Since the above studies showed that gene *dhc* is dispensable for dicamba demethylation and that 5,10-CH_2_-THF is unstable and rapidly spontaneously transforms to THF, we hypothesized that MthfD alone could regenerate THF from 5-CH_3_-THF during dicamba demethylation. To confirm this hypothesis, an enzymatic reaction was designed. The reaction mixture contained 1.0 mM dicamba, 0.1 mM THF (or 0.1 mM 5-CH_3_-THF), 2.0 mM menadione, 0.01 mg of purified Dmt and 0.01 mg of purified MthfD. Theoretically, the conversion of 1.0 mM dicamba requires 1.0 mM THF and yields 1.0 mM dichlorosalicylic acid (3,6-DCSA) and 1.0 mM 5-CH_3_-THF. The enzymatic assay showed that although only 0.1 mM THF was added, the 1.0 mM dicamba was completely transformed after incubation for 10 min, and no 5-CH_3_-THF accumulation was observed when the mixture contained both Dmt and MthfD (Fig. S4AII).

Simultaneously, only approximately 5% dicamba was transformed when the mixture contained only Dmt (Fig. S4AIII). When 0.1 mM 5-CH_3_-THF was added to the reaction mixture, a similar result was obtained: 95% of the added 1.0 mM dicamba was transformed in the mixture containing both Dmt and MthfD (Fig. S4BII), whereas the dicamba was not degraded in the reaction mixture containing only Dmt (Fig. S4BIII). The results indicated that THF could be continuously regenerated from 5-CH_3_-THF by MthfD during dicamba demethylation *in vitro*.

## DISCUSSION

BLAST analysis in the GenBank database indicated that MthfD is most related to MetF. However, MthfD shares very low identity with characterized MetFs, e.g., only 26% identity with *E. coli* MetF. The phylogenetic relationship between MthfD and characterized MetFs is also very distant. In the phylogenetic tree of MthfD and some related MetFs (Fig. 6), there are obviously two distinct branches. One branch consists of MetFs from *E. coli*, yeast, *Caenorhabditis elegans*, *Arabidopsis thaliana* and human ect., the physiological function of these MetFs is reported to be responsible for reduction of 5,10-CH_2_-THF to 5-CH_3_-THF (24–27, 36). Another branch is composed of four MthfDs/MetFs from *R. dicambivorans* Ndbn-20, *M. chloromethanicum* CM4 and *S. paucimobilis* SYK-6. All the genes coding these MthfDs/MetFs are located in THF-dependent demethylase gene clusters, and their roles are predicted as 5-CH_3_-THF dehydrogenase (6, 16, 18, 21).

**FIG 6.**
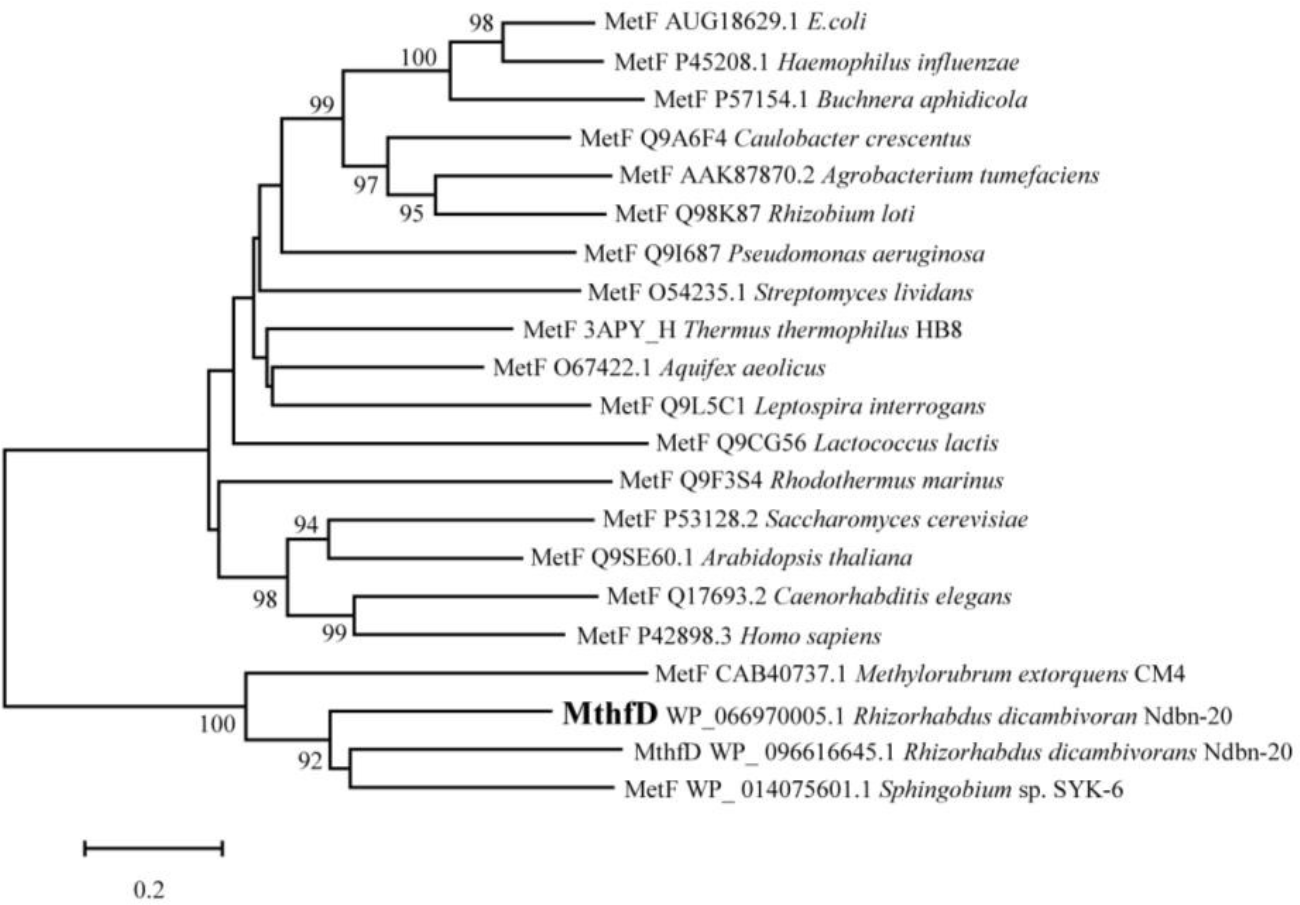
Phylogenetic tree constructed based on the alignment of MthfD with related MetFs. The multiple-alignment analysis was performed with ClustalX v2.0, the phylogenetic tree was constructed by the neighbor-joining (NJ) method using MEGA 5.0, bootstrap values (based on 1,000 replications) are indicated at branch nodes. Bar, 0.2 substitution per nucleotide position. Each item is arranged in the following order: protein name, GI number and organism name.

Furthermore, MthfD differs from *E. coli* MetF in structure, catalytic activities and characteristics. Mature MthfD is a monomer that is different from the MetFs of *E. coli* (tetramer) (26) and *M. thermoacetica* (hexamer) (29). MthfD was fairly stable and tightly bound with flavin cofactor, whereas *E. coli* MetF was fragile and readily lost its activity and flavin cofactor. Both MthfD and *E. coli* MetF exhibited 5-CH_3_-THF dehydrogenase activity, however, the *K*_m_ of MthfD was only half of that of *E. coli* MetF, and the *K*_cat_/*K*_m_ was 2.7 times higher than that of *E. coli* MetF, indicating that MetF had higher affinity and catalytic efficiency than that of *E. coli* MetF. Furthermore, an unexpected result was that MthfD could not catalyze the reverse reaction, or its 5,10-CH_2_-THF reductase activity was too low to be detected; in contrast, the *E. coli* MetF displayed 5,10-CH_2_-THF reductase activity as expected, indicating that reaction catalyzed by *E. coli* MetF is reversible. 5,10-CH_2_-THF reductase is a very important enzyme in organisms, its catalytic product 5-CH_3_-THF is used to synthesize methionine which is a methyl donor for methylation of DNA, RNA, protein and lipid and synthesis of other amino acids such as cysteine and taurine (22, 24). When *R. dicambivorans* Ndbn-20 cells grown on dicamba, 5-CH_3_-THF was produced from dicamba demethylation, when grown on other carbon source, 5-CH_3_-THF should be produced from 5,10-CH_2_-THF under the catalysis of an unknown 5,10-CH_2_-THF reductase.

The fact that MthfD exhibited 5-CH_3_-THF dehydrogenase activity and no 5,10-CH_2_-THF reductase activity suggests that the physiological function of MthfD is to catalyze the dehydrogenation of 5-CH_3_-THF to 5,10-CH_2_-THF during dicamba demethylation in *R. dicambivorans* Ndbn-20. RT-PCR results showed that *mthfD* was transcribed in dicamba-cultured cells, which was consistent with our previous research (16), while the homologous gene of *mthfD* in cluster scaffold 50 was not transcribed. Furthermore, the *mthfD*-disrupted mutant Ndbn-20∆*mthfD* lost the ability to degrade and utilize dicamba, although the mutant possessed an intact dicamba methyltransferase gene *dmt*. Enzymatic study also showed that the cell lysate of mutant Ndbn-20∆*mthfD* lost 5-CH_3_-THF dehydrogenase activity. This evidences further demonstrated that MthfD physiologically acts as a 5-CH_3_-THF dehydrogenase.

According to a previously proposed pathway for THF regeneration from 5-CH_3_-THF produced in substrate demethylation, Dhc is a necessary bifunctional enzyme that is responsible for the dehydrogenation of 5,10-CH_2_-THF to 5,10-CH=THF and subsequent cyclohydrolysis of 5,10-CH=THF to 10-CHO-THF (6, 18, 21). However, it is interesting that the *dhc*-disrupted mutant Ndbn-20∆*dhc* could still degrade and utilize dicamba with a rate similar to that of the wild-type. There are two possible explanations for this phenomenon: one possibility is that the two reactions from 5,10-CH_2_-THF to 10-CHO-THF are performed by other isozyme since bioinformatic analysis showed that there were three other *dhc*-like genes in the genome of *R. dicambivorans* Ndbn-20 besides the *dhc* in scaffold 66. Another possibility is that the 5-CH_3_-THF produced during dicamba demethylation is transformed by MthfD to 5,10-CH_2_-THF, which is spontaneously converted to THF via a nonenzymatic reaction (Fig. 1C), only MthfD is required, and Dhc is likely not involved in this reaction. This hypothesis was supported by the fact that the addition of a small amount of THF is sufficient for conversion of a large amount of dicamba in enzymatic reaction mixture containing both Dmt and MthfD and that the cell lysate of mutant Ndbn-20∆*dhc* displayed the wild-level of convention rate of 5-CH_3_-THF to THF.

Similar gene disruption results have also been reported in the vanillate-utilizing strain *S. paucimobilis* SYK-6 and chloromethane-utilizing strain *M. chloromethanicum* CM4. In both strains, substrate demethylation was catalyzed by a THF-dependent methyltransferase system. The *metF*-disrupted mutants of *S. paucimobilis* SYK-6 and *M. chloromethanicum* CM4 lost the ability to grow on vanillate or chloromethane, and the cell lysates of these *metF*-disrupted mutants lost the 5-CH_3_-THF dehydrogenase activities (17, 18, 21). The *folD* (*dhc*)-disrupted mutant of *M. chloromethanicum* CM4 retained the ability to grow on chloromethane with a growth yield similar to that of the wild-type (21). All these results showed that that *metF* is essential and *dhc* is dispensable for the THF-dependent demethylase system in these strains. However, the location of *dhc* near and co-transcription with the THF-dependent demethylase gene support the idea that *dhc* is involved in THF metabolism. Further study is needed to elucidate the THF regeneration mechanism and the physiological function of Dhc.

## MATERIALS AND METHODS

### Chemicals and media

Dicamba (99.3% purity), THF (⩾ 90% purity), and 5-CH_3_-THF (97% purity) were purchased from Sigma-Aldrich (Shanghai, China). All other chemicals and reagents (>98% purity) were purchased from Sangon Biotech Co., Ltd. (Shanghai, China). LB broth was purchased from Difco Laboratories (Detroit, MI, USA). LB (1/5) was prepared by diluting LB broth 4-fold with distilled water, and 2% agar was added for solid plate. Minimal salt medium (MSM) was prepared as described by Yao et al. 2016 (16).

### Bacterial strains, plasmids and culture conditions

The bacterial strains and plasmids used in this study are listed in Table 1. The primers are listed in Table 2. *R. dicambivorans* Ndbn-20 and its mutants were cultured in 1/5 LB broth or MSM supplemented with 1.0 mM dicamba at 30°C, and all *E. coli* strains were grown aerobically at 37°C. Antibiotics were added at the following concentrations: streptomycin, 100 μg/mL; kanamycin, 100 μg/mL; chloramphenicol, 30 μg/mL; gentamicin, 50 μg/mL for *E. coli* strains and 15 μg/mL for *R. dicambivorans* Ndbn-20 strains.

**TABLE 1.**
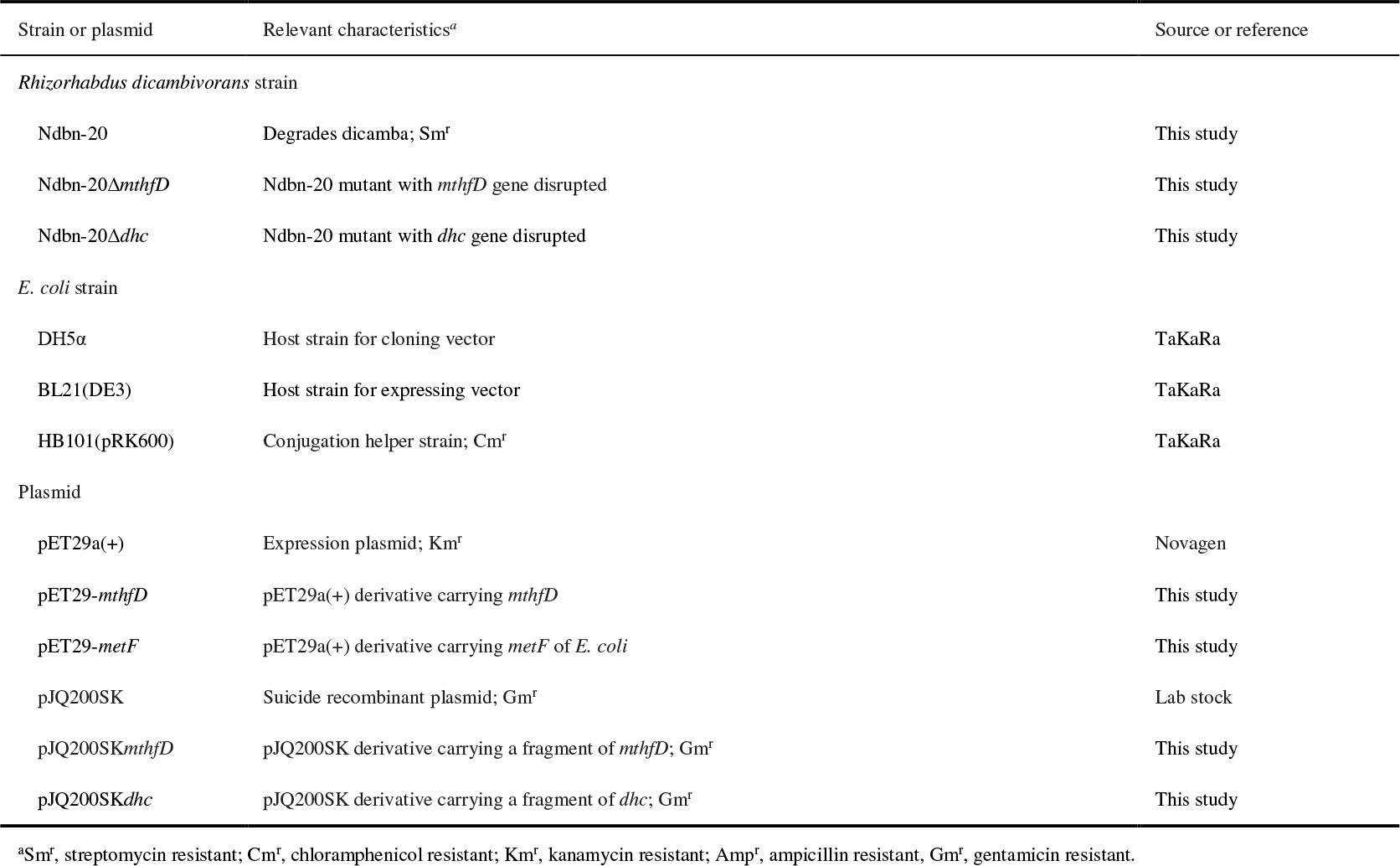
Strains and plasmids used in this study

**TABLE 2.**
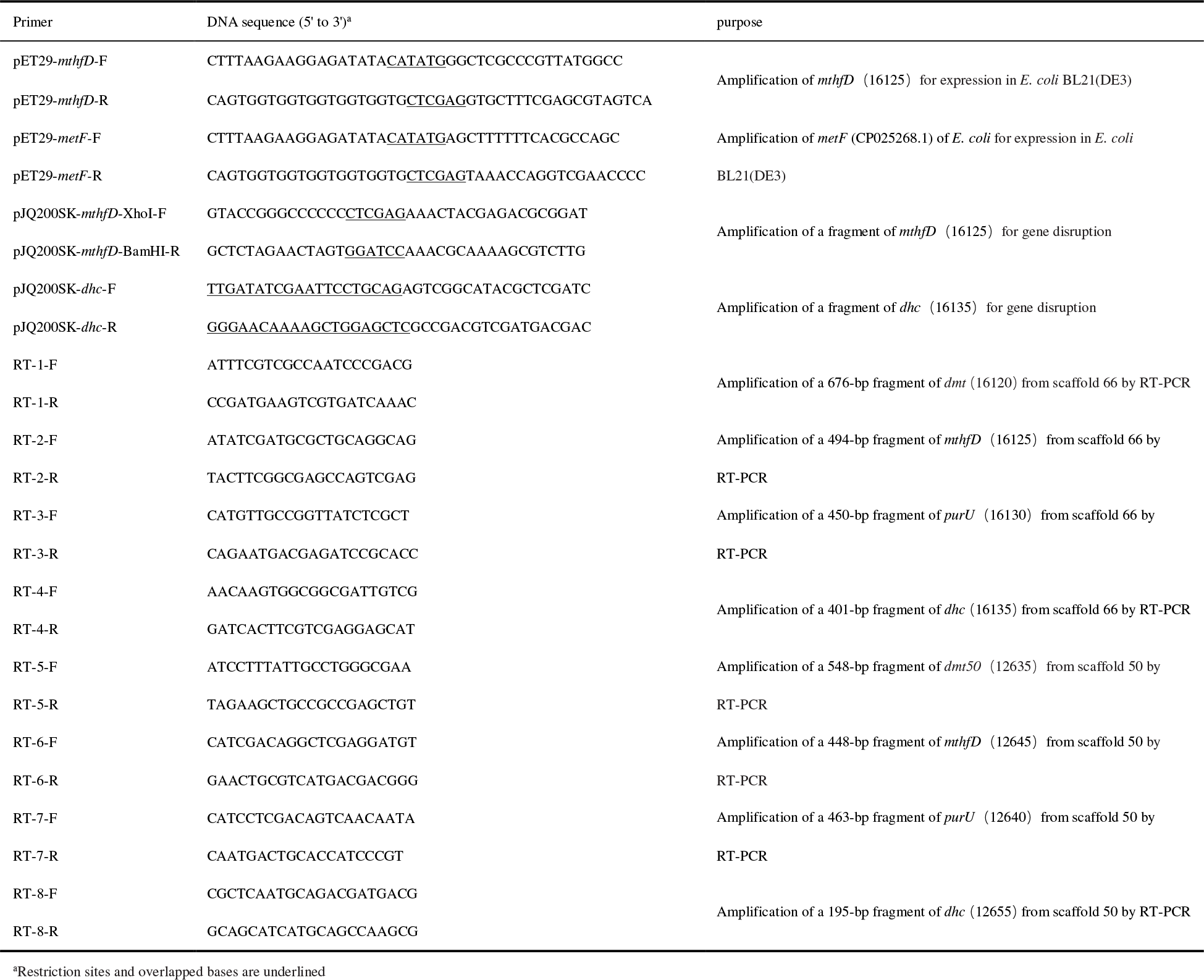
PCR primers used in this study

### Heterogenous expression of *mthfD/metF* and purification of the His_6_-tagged proteins

The sequence of *mthfD* was amplified by PCR using 2×Phanta Master Mix (Vazyme Biotech Co., Ltd) using the primer pairs pET29-*mthfD*-F and pET29-*mthfD*-R and the Ndbn-20 genomic DNA as a template. The PCR product was cloned into the NdeI-XhoI site of pET29a(+) using the ClonExpress^®^ II One Step Cloning Kit (Vazyme Biotech Co., Ltd.) to generate the recombinant plasmid pET29-*mthfD*. Then, pET29-*mthfD* was transformed into *E. coli* BL21 (DE3). The transformed cells were grown in LB broth containing 100 μg/mL kanamycin at 37°C until the turbidity of the culture at 600 nm (OD_600 nm_) reached 0.6. Then, the expression of *mthfD* was induced by adding 0.4 mM isopropyl-β-D-thiogalactopyranoside (IPTG), and the cells were grown for another 12 h. The cells were then harvested by centrifugation at 12,000 × g for 1 min, resuspended in ice-cold 20 mM Tris-HCl buffer (pH 8.0), and disrupted by sonication. The cell lysate was centrifuged at 12000 × g for 30 min at 4°C, and the supernatant was obtained. The C-terminal His_6_-tagged protein was purified from the supernatant using Co^2+^-affinity chromatography (His GraviTrap™ TALON^®^, GE Healthcare Life Sciences Co., Ltd.) following the manufacturer’s instructions. The resultant fraction was dialyzed with 20 mM Tris-HCl buffer at 4°C for 24 h. The molecular weight of the purified protein was determined by 4-20% SDS-PAGE and Coomassie brilliant blue staining. The protein concentration was quantified by the BCA method using bovine serum albumin as a standard (BCA Protein Assay Kit, Sangon Biotech Shanghai Co., Ltd.). The molecular mass of the native protein was determined by gel filtration (32). Briefly, the purified protein was loaded onto Superdex 2000 column (GE Healthcare AKTAprime liquid chromatography system) pre-equilibrated with 20 mM Tris-HCl buffer. The column was washed with the same buffer at a flow rate of 0.4 mL/min. The standard proteins used were myosin (200.0 kDa), β-galactosidase (116.0 kDa), phosphorylase b (97.2 kDa), bovine serum albumin (66.4 kDa) and ovalbumin (44.3 kDa). The *metF* gene of *E. coli* was also expressed using pET29a(+) and the His_6_-tagged MetF was purified according to the above method.

### Enzyme activity assay

The 5-CH_3_-THF dehydrogenase activity was determined using HPLC. A 1-mL reaction mixture containing 5-CH_3_-THF (10 μM to 300 μM), 0.5 mM menadione and 20 mM Tris-HCl buffer was pre-incubated at 30°C for 5 min, then an appropriate amount of enzyme was added to initiate the reaction. After incubation at 30°C for 10 min, the reaction was terminated by boiling at 100°C for 1 min. The conversion of 5-CH_3_-THF was analyzed by HPLC as described below. One unit of dehydrogenase activity was defined as the amount of enzyme required to catalyze the conversion of 1 μmol of 5-CH_3_-THF per minute. The *K*_m_ and *K*_cat_ were measured independently three times. The kinetic values were calculated via nonlinear regression (33) fitting to the Michaelis-Menten equation using OriginPro 9.0 (OriginLab Corporation, Northampton, MA).

The 5,10-CH_2_-THF reductase activity of the enzyme was determined using a UV2450 spectrophotometer (Shimadzu). 5,10-CH_2_-THF was prepared via nonenzymatic reaction of THF and formaldehyde as described in a previous study (34). After preincubation of the assay buffer (1.0 mL of 20 mM Tris-HCl) in a cuvette at 30°C, 0.5 mM NADH and 0.01 mg of purified MthfD or *E. coli* MetF were added into the cuvette, and the reaction was monitored at 340 nm every 30 s.

### RNA preparation and transcriptional analysis

Cells of strain Ndbn-20 were cultivated in MSM supplemented with 1.0 mM dicamba. The cells were harvested until approximately 60% of the added dicamba was transformed. Total RNA was isolated with the RNA Isolation Kit (TaKaRa). To remove any genomic DNA contamination, the recovered RNA was treated with gDNA Eraser (TaKaRa) at 42°C for 2 min. RT-PCR was performed with the PrimeScript RT Reagent Kit (TaKaRa) and the primers listed in Table 2. All samples were analyzed in triplicate.

### Disruption of *mthfD* and *dhc* in *R. dicambivorans* Ndbn-20 by homologous recombination

The knockout plasmids pJQ200SK-*mthfD* and pJQ200SK-*dhc* were constructed by fusing a middle fragment (approximately 500 bp) of the target gene to the PCR-linearized suicide plasmid pJQ200SK (35) using the One-step Cloning Kit (Vazyme Biotech Co., Ltd.). The homogenous fragments were amplified with the primers listed in Table 2. The recombinant plasmids were transformed into *E. coli* DH5α and then introduced into *R. dicambivorans* Ndbn-20 by triparental mating using pRK600 as a helper. The candidate mutants were screened on 1/5 LB agar containing 15 μg/mL gentamicin and 100 μg/mL streptomycin. The disruption events in the mutants were verified by PCR and sequencing. The abilities of the mutants to degrade dicamba were determined via a whole-cell biotransformation test in MSM as described by Li et al. (35). The initial concentration of dicamba was 1.0 mM, and the conversion of dicamba was monitored by HPLC as described below.

To investigate the conversion of 5-CH_3_-THF by the cell lysates of the mutants and wild-type, dicamba-cultured cells were harvested in the logarithmic phase and disrupted by sonication. Then the crude cell lysate was centrifuged at 12,000 × g for 30 min at 4°C to remove cell debris and undisrupted cells. The supernatant was obtained as the cell lysate. A 300-μL enzymatic mixture containing 1.0 mM 5-CH_3_-THF, 1.0 mM menadione, 200 μL of cell lysate and 20 mM Tris-HCl buffer was incubated at 30°C for 10 min, and the conversion of 5-CH_3_-THF was determined by HPLC as described below. To study the dicamba methyltransferase activity, 300 μL of enzymatic mixture containing 1.0 mM dicamba, 1.0 mM THF, 200 μL of cell lysate and 20 mM Tris-HCl buffer was incubated at 30°C for 1 h, and the conversion of dicamba was determined by HPLC as described below.

### Analytical methods

The culture or enzymatic sample was centrifuged at 12,000 × g for 5 min and then filtered through a 0.22-μm Millipore membrane to remove the cell or protein precipitate. An UltiMate 3000 Titanium system (Thermo Fisher Scientific) equipped with a C_18_ reversed-phase column (4.6 × 250 mm, 5 μm; Agilent Technologies) was used for HPLC analysis. For detection of dicamba, the mobile phase was a mixture of ultrapure water, acetonitrile, methanol and acetic acid (58.4:31.7:7.5:2.4, vol/vol/vol/vol) at a flow rate of 1.0 mL/min; the column temperature was set as 30°C, the injection volume was 20 μL, the detection wavelengths for dicamba and 3,6-DCSA were 275 nm and 319 nm, respectively. Detection of THF and 5-CH_3_-THF was performed according to the method described by Chen et al. (16, 36, 37). The sample was dissolved in isovolumetric 0.1 M KH_2_PO_4_ buffer added with 1% ascorbic acid and 0.1% β-mercaptoethanol (pH 6.8), the mobile phase was a mixture of 0.05 M KH_2_PO_4_ buffer (pH 3.0) (90%) and acetonitrile (10%), the flow rate was 0.8 mL/min, the detection wavelength for both THF and 5-CH_3_-THF was 315 nm. All assays were performed as three independent experiments. The formation of formaldehyde was identified by HPLC after derivatization with 2,4-dinitrophenylhydrazine. A 500-μL sample was mixed with isovolumetric 0.6 g/L 2,4-dinitrophenylhydrazine, the mixture was incubated at 60°C for 1 h and then cooled to room temperature. The HPLC conditions were as follows: the mobile phase was a mixture of methanol and water (70:30, vol/vol) at a flow rate of 1.0 mL/min, the column temperature was set at 40°C, the injection volume was 20 μL, the detection wavelength was 365 nm.

## Accession number

The chromosomal genome of *R. dicambivorans* Ndbn-20 has been deposited in the GenBank database under the accession number CP023449, the GenBank accession number of MetF of *E. coli* was CP025268.1.

## Acknowledgments

This work was supported by the National Natural Science Foundation of China (No. 31770117, 31570105) and Science and Technology Project of Jiangsu province (No. BE2016374).

